# Effects of Spearman’s and Pearson’s correlations on construction of cancer regulatory networks and biomarker selection

**DOI:** 10.1101/2025.10.19.683264

**Authors:** Houri Razavi, Amirhosein Javadpour

## Abstract

Correlation methods play an important role in machine learning. They apply as a similarity or distance measure in machine learning models such as recommender systems, clustering methods, and distance-based algorithms such as KNN, leading to more accurate and interpretable results. Pearson’s correlation is one of the most common methods for measuring dependence between biological variables. However, it works well only when data are linearly associated and follow a normal distribution. In cancer, most molecules do not follow a normal distribution. A sudden increase or decrease of gene expression in high stages of cancer usually leads to exponential distribution. In such circumstances, rank-based correlations would be a better choice for association detection. Misusing the Pearson correlation coefficient in cases where data exhibit a non-linear relationship can lead to misleading results and incorrect interpretations such as the exclusion of important features and selection of inappropriate relationships.

In this paper, we applied four correlation metrics to define the association between genes in cancer networks. There was a significant difference between the rate of survival-related hubs among Pearson’s and Spearman’s based networks. For instance, in BLCA, the rate of survival-related hubs was 81% for the Spearman-based network vs 68% for the Pearson-based network. This discrepancy was 53 % vs 42 % for KIRC. Furthermore, Spearman-based networks enriched more GO terms, and their cancer-related enriched GO terms were twice as many as those of Pearson’s. The source code of the study is available at https://github.com/Amirhosein-JVD/Correlation-Analysis

## Introduction

Biological characteristics can arise from complex interactions between the cell’s molecules, such as DNAs, RNAs, and proteins [1]. Understanding the structure and the dynamic of biological networks contributes not only to identifying potential biomarkers and drug targets but also allows us to characterize networks and sub-networks of biomolecular interactions critical for the emergence of a disease state [2].

Biological network analysis has largely benefited from machine learning in identifying network architectures. A remarkable example is learning patterns from input data to generate biologically relevant gene regulatory networks, with interesting applications in the identification of drivers of drug response or disease phenotypes [2].

Correlation calculation is a common technique used in preprocessing steps before applying machine learning methods. For example, you can apply correlation calculation as a feature selection method before applying the machine learning model. Furthermore, Correlation is a measure of similarity that is applied as a part of machine learning methods. Unsupervised learning such as clustering methods uses correlation among biomolecules as a metric to define the distance between two objects (e.g. two genes) [3,4,5,6,7].

In biological networks, each network node is a biological molecule and, in many cases, the link between nodes is defined by correlation. Pearson’s correlation is the most common method used to measure dependence between molecules to link biological nodes, but it works well only when data are linearly associated [5,6]. However, empirical research in molecular biology shows these associations are not necessarily linear. To identify associations not limited to linear associations, but dependent in a broad sense, several methods have been expanded. Most of these methods are based on rank correlation and information theory, but are less used in applications, most probably because of a lack of clear guidelines for their use [4,6]. For instance, Spearman’s rank correlation assesses how well the correlation between two variables can be formulated by a monotonic function [8,9]. Furthermore, linear and monotonic dependence are not the only ones observed in a real biological system, many other complex relationships are also observed in biological Systems. Consequently, if the correlation method is limited to linear dependence measures in the biological network construction, the network’s ability to model the accurate biological structure will also be limited.

This study compares the popular correlation methods (Pearson’s, Spearman’s, Kendall’s, and distance correlations) used to discover dependency between network nodes, especially based on gene expression profiles. The methods were applied to construct gene co-expression networks in Stomach (STAD), Colon (COAD), Bladder (BLCA), and Kidney (KIRC) cancers. High-degree (hub) nodes of these networks were selected for further analysis. Finally, the results proved that Spearman correlation was more successful in extracting nonlinear associations that led to influential pivotal gene selection. These genes are suitable potential cancer biomarkers that are beneficial in cancer diagnosis or drug discovery.

## Results

### Data distribution

Pearson’s correlation is a measure of strength in linear correlation between variables, but it works well when the variables have normal distribution [10,11,12,13]. In biological networks, molecule expressions usually do not follow the normal distribution, especially when you study those molecules that are the disease biomarkers [4]. In our study, gene expression profiles were log-transformed to achieve a normal distribution. However, the majority of genes did not exhibit normality even after transformation. To formally assess the distribution of network genes, we applied the Shapiro-Wilk test [14], which calculates a p-value for each gene. A p-value greater than 0.05 indicates that the gene’s expression follows a normal distribution. Notably, none of the initial nodes in our cancer gene networks had p-values exceeding this threshold, indicating that none of these genes were normally distributed.

In most cases, the gene expression patterns were strongly non-linear and followed an exponential distribution. This outcome is likely due to the initial selection of genes with significantly altered expression levels, which tend to show abrupt increases or decreases, characteristics that typically lead to an exponential distribution.

Fig 1 depicts normal quantile-quantile (QQ) plots of the four gene expression profiles obtained from the cancer datasets. A QQ plot compares a gene expression profile on the vertical axis to a standard normal population on the horizontal axis. If the data were normally distributed, the points would align along a straight line. However, the plots deviate significantly from linearity, further confirming the non-normal nature of the data. Given these findings, Pearson’s correlation application, which assumes normality of data distribution, is not appropriate in this context despite its widespread use.

**Fig 1.**
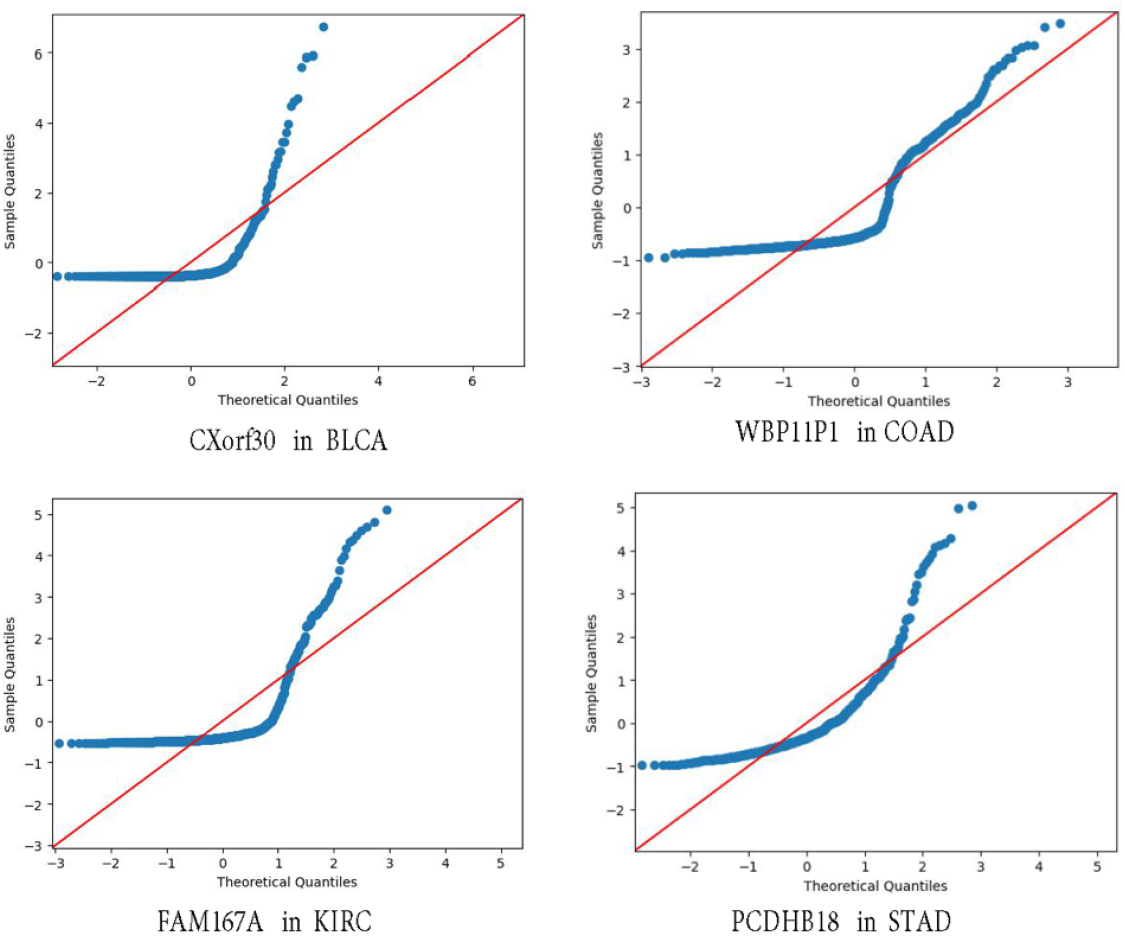
Normal QQ plots of gene expression profiles obtained from cancer datasets. The gene expression profiles do not follow a normal distribution and exhibit features of exponential distributions.

### Spearman’s correlation better suits cancer regulatory networks

Associations between biological molecules in biological networks are usually complex. This complexity intensifies in the case of studying cancer-related networks because of the abnormal cell function. Fig 2 provides several examples of gene expression relationships observed between pairs of genes in cancer datasets. Each subplot presents the Pearson, Spearman, Kendall, and Distance Correlation coefficients for a pair of genes. The concentration of most points in the lower-left corner indicates that gene expression levels are similar across the majority of samples, with notable deviations observed in a small number of samples, typically those in more advanced stages of cancer. Number of samples in each subplot corresponds to the number of tumor samples in each cancer datasets. This number is 414 in BLCA, 482 in COAD, 541 in KIRC, and 415 in STAD datasets.

**Fig 2.**
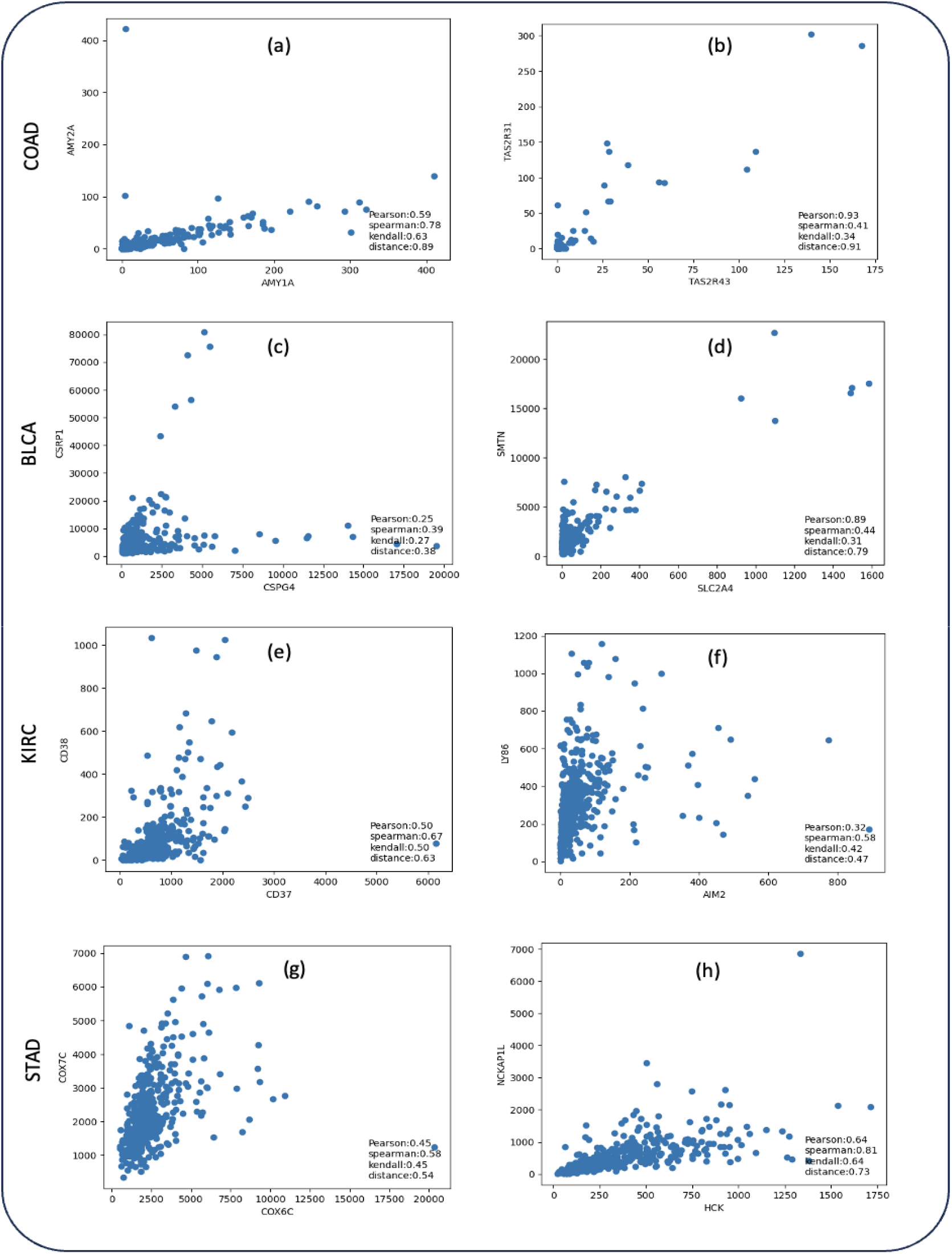
Examples of gene expression associations observed between gene pairs in cancer datasets. The horizontal and vertical axes each represent the expression level of a gene across different cancer samples. Each point corresponds to the expression levels of two genes in a single sample.

A small difference between the Pearson coefficient and the others may indicate a linear relationship. A higher Pearson coefficient compared to Spearman’s suggests the presence of outliers within a linear association. Conversely, a lower Pearson coefficient relative to Spearman’s implies a non-linear relationship or the influence of outliers in a non-linear context. These patterns were also detected in [4].

In Fig 2 (b) and (d), Pearson’s correlation is noticeably higher than Spearman’s reflecting the impact of outliers on a linear association. In Fig 2 (e), (f), and (g) Spearman’s correlation is higher due to non-linear relationships. The genes shown in the subplots are part of the initial set of network genes, selected based on their differential expression between tumor and normal samples.

In this study, as in many others, the objective is to identify potential cancer biomarkers based on gene expression data. Removing samples considered outliers is not an option, as these samples often exhibit the most significant biological changes. Therefore, using Spearman’s correlation instead of Pearson’s is a more suitable choice, as it is more robust to outliers and can also capture non-linear associations.

After calculating correlation values for gene pairs, we observed that in some cases, Pearson’s correlation was higher than Spearman’s, and in others, the reverse held true. This pattern is also evident in Fig 2.

Fig 3 illustrates the proportion of gene pairs with relatively similar correlation values (difference < 0.05) between Pearson and Spearman, compared to gene pairs with substantial divergence between the two measures. In the BLCA dataset, 40.63% of gene pairs showed a higher Spearman correlation, indicating that most gene associations follow a non-linear pattern. In the KIRC dataset, the difference is even more pronounced, with 18.65% of gene pairs showing Pearson > Spearman and 44.08% showing Spearman > Pearson. This greater disparity is likely due to the lower number of outliers and the prevalence of non-linear associations in KIRC. These observations are consistent with the patterns shown in Figs 2(e) and 2(f). Overall, the results confirm that using Spearman’s correlation instead of Pearson’s yields more accurate and biologically meaningful insights in cancer gene expression analysis.

**Fig 3.**
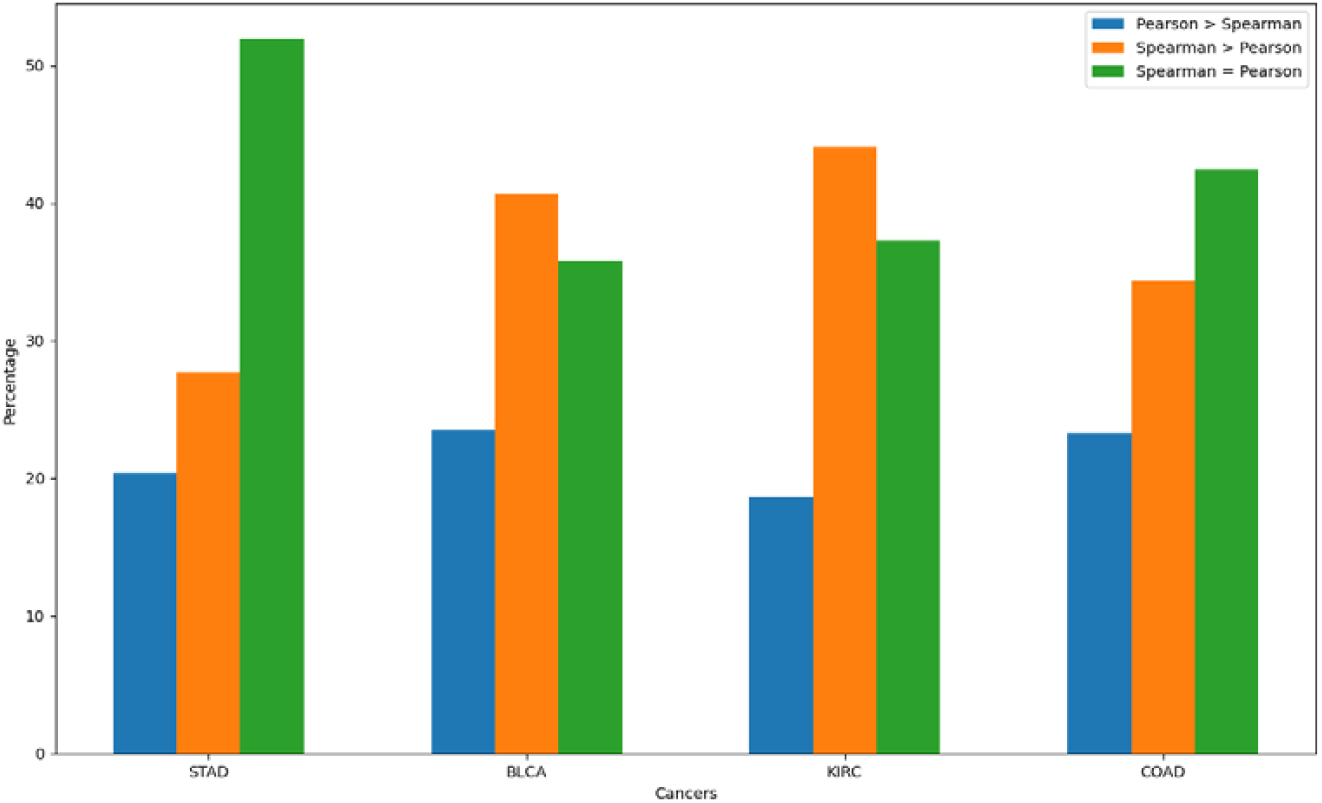
Comparative analysis of correlation discrepancies across various cancer types. The horizontal axis represents the cancer types, and the vertical axis shows the percentage of gene pairs. The green bars represent the percentage of gene pairs with similar Pearson and Spearman correlation values, while the blue and orange bars indicate gene pairs with significant discrepancies between the two.

All correlation computation were performed on a desktop computer running macOS, equipped with an M1 chip, an 8-core GPU, and 16 GB of RAM. Table 1 presents the computation time required to calculate correlations between the initial genes and all other genes in each cancer network.

**Table 1.** Computation Time for Correlation Values Across Cancer Networks.

|  | *Pearson* | *Spearman* | *Kendall* | *Distance* | *Num_genes* |
| --- | --- | --- | --- | --- | --- |
| *STAD* | 0:00:01.164018 | 0:00:00.639223 | 0:02:48.679975 | 0:06:46.528497 | 1511 |
| *BLCA* | 0:00:03.919316 | 0:00:02.084337 | 0:09:34.952293 | 0:20:16.700476 | 2768 |
| *KIRC* | 0:00:03.304213 | 0:00:01.783959 | 0:06:46.757547 | 0:17:37.895606 | 2222 |
| *COAD* | 0:00:03.136705 | 0:00:01.689368 | 0:06:57.320534 | 0:15:57.668590 | 2303 |

The number of genes is consistent across the networks. Pearson and Spearman correlations were both computed rapidly, with Spearman demonstrating slightly better efficiency. Although Kendall and Distance correlations occasionally yielded efficient results, their computational complexity is significantly higher and not comparable to the speed and scalability of Spearman and Pearson methods.

### Effects of correlation metrics in survival-related biomarker selection

Correlation metrics were employed to construct cancer gene networks. Initially, genes exhibiting significant differential expression in each cancer dataset were selected as the initial network nodes (Supplementary File 1). Next, the correlation metrics were used to establish associations between these genes, forming the network structure. *Finally, the high-degree nodes were considered as potential cancer biomarkers. We select the top 100 genes as network hubs (any other number, like 50, does not affect the results). Additionally, genes whose expression levels were associated with patients survival status (alive or deceased) were identified as survival-related genes. This analysis was performed using Kaplan–Meier survival curves and log-rank tests based on gene expression profiles and corresponding survival data. (Supplementary File 2)*. Genes with the highest degrees (i.e., most connections) were then identified as potential cancer biomarkers. The top 100 genes were selected as network hubs; using a different cutoff (e.g., 50) did not significantly alter the results. Additionally, survival-associated genes were identified based on their expression profiles using Kaplan–Meier analysis and log-rank tests (Supplementary File 2).

Finally, we examined the overlap between network hubs and survival-related genes within each network to identify the survival-related hubs and determine their proportion among the total hub genes (Table 2). As shown in Table 2, 81 % of hub genes in the BLCA Spearman-based network were related to overall survival, compared to 68 % in BLCA Pearson-based. These results demonstrate that Spearman’s correlation is capable of effectively detecting both linear and non-linear associations. This finding is consistent with the results presented in Fig 3, where 40.63% of gene pairs in the BLCA dataset exhibited higher Spearman correlation values compared to Pearson correlation. This observation suggests a high prevalence of non-linear relationships among gene pairs in BLCA.

**Table 2.**
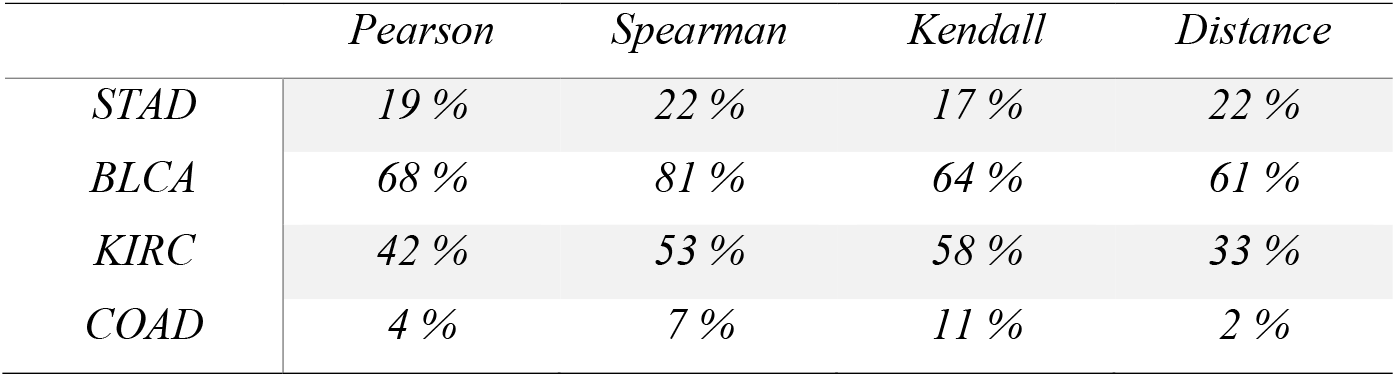
Proportion of Hub Genes Associated with Overall Survival Across Networks.

|  | *Pearson* | *Spearman* | *Kendall* | *Distance* |
| --- | --- | --- | --- | --- |
| *STAD* | 19 % | 22 % | 17 % | 22 % |
| *BLCA* | 68 % | 81 % | 64 % | 61 % |
| *KIRC* | 42 % | 53 % | 58 % | 33 % |
| *COAD* | 4 % | 7 % | 11 % | 2 % |

Similar results were observed in the KIRC and COAD networks. The relatively low proportion of overall survival-related hub genes in COAD can be attributed to the high survival rate within this cancer cohort. In contrast, the STAD network exhibited less variation among the four correlation-based networks, which is likely due to the higher proportion of linear associations (51.9%) among gene pairs, as shown in Fig 3. Collectively, these findings further support the superiority of Spearman correlation in the analysis of cancer gene regulatory networks.

To evaluate the efficiency of different correlation metrics in identifying survival-related hub genes, we analyzed the overlap of survival-related hubs across correlation-based networks for each cancer type, as depicted in Fig 4. In the case of BLCA, the networks constructed using Spearman’s and Kendall’s correlations shared 52 common survival-related hubs, indicating the highest degree of concordance among the BLCA networks. In contrast, the overlap between the Pearson- and Spearman-based networks included 34 common survival-related hubs, suggesting that approximately 41% of the survival-related hubs identified by Spearman’s method were also captured by Pearson’s correlation. These findings support the observations presented in Fig 3 and further highlight the superior ability of Spearman’s correlation to detect both linear and non-linear associations relevant to patient survival.

**Fig 4.**
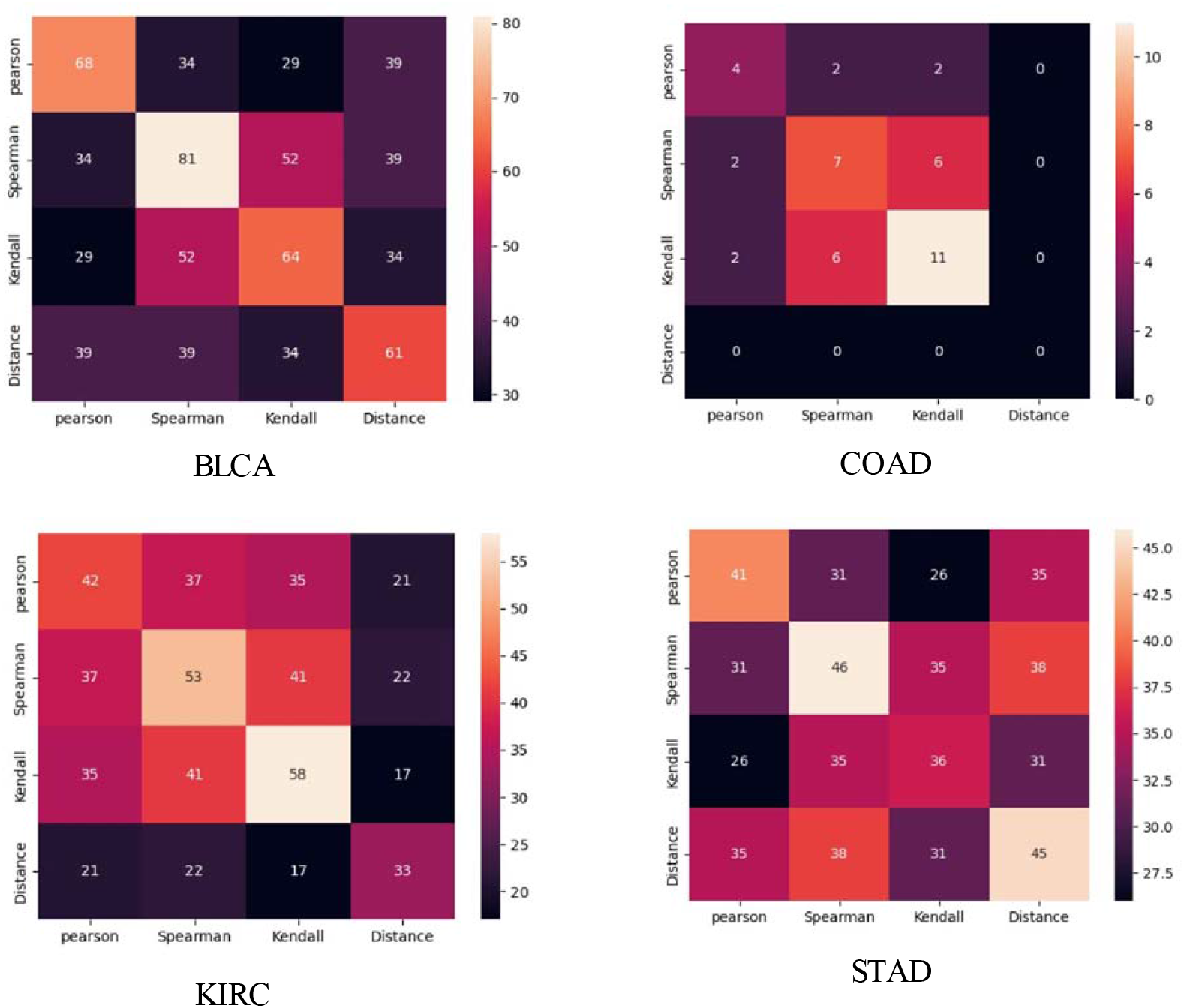
Heatmap depicting overlap of survival-related hub genes across four correlation-based networks in each cancer type.

### Effects of correlation metrics on Enriched GO terms and pathways

To explore the biological functions and pathways associated with hub genes in each cancer network, enrichment analysis was conducted using g:Profiler for the top 100 high-degree genes derived from both Pearson and Spearman-based networks (Supplementary File 3). Across all four cancer datasets, the number of enriched Gene Ontology (GO) terms and pathways identified in Spearman-based networks was significantly greater than those identified in Pearson-based networks, as shown in Table 3. This observed difference may be attributed to Spearman’s ability to capture strong associations between gene expression levels when a genuine functional relationship exists. In other words, the results suggest that Spearman’s correlation is more effective than Pearson’s in detecting functionally relevant gene-gene associations, thereby providing a richer and more biologically meaningful network structure.

**Table 3.** Enrichment Analysis Results for Each Cancer Network. The table summarizes the Gene Ontology (GO) enrichment analysis of the top 100 high-degree hub genes identified in each cancer network. Enrichment results are categorized into three main GO domains: Molecular Function (MF), Biological Process (BP), and Cellular Component (CC).

| Cancer Network | GO:MF | GO:BP | GO:CC | KEGG | REACTOM | Cancer-related GO terms |
| --- | --- | --- | --- | --- | --- | --- |
| BLCA - Pearson | 9 | 53 | 32 | 11 | 9 | 2 |
| BLCA - Spearman | 12 | 59 | 23 | 2 | 4 | 14 |
| COAD - Pearson | 1 | 0 | 0 | 2 | 3 | 0 |
| COAD - Spearman | 2 | 14 | 9 | 3 | 6 | 0 |
| KIRC - Pearson | 9 | 160 | 12 | 10 | 8 | 12 |
| KIRC - Spearman | 12 | 330 | 40 | 15 | 22 | 25 |
| STAD - Pearson | 0 | 1 | 3 | 0 | 0 | 0 |
| STAD - Spearman | 0 | 4 | 5 | 0 | 0 | 1 |

To assess the quality of enriched GO terms, we focused specifically on those associated with cancer. Chen et al. [15] conducted a comprehensive review compiling studies that mapped Gene Ontology (GO) terms to cancer hallmarks. Based on their findings, we aggregated all reported associations from the reviewed studies that linked GO terms to cancer hallmarks (Supplementary File 4). For instance, in the Spearman-based BLCA network, 59 biological processes were enriched, among which 9 were linked to the hallmark “Activating Invasion and Metastasis,” 3 to “Inducing Angiogenesis,” and 2 to “Tumor-Promoting Inflammation.” Overall, our results demonstrated that the number of cancer-related enriched GO terms in Spearman-based networks was consistently twice as high as in Pearson-based networks across all cancer types (Table 3). Detailed gene set enrichment results and their alignment with cancer hallmarks are provided in Supplementary File 1.

## Discussion

*Cancer is rarely a consequence of an abnormality in a single gene*; instead, it reflects disturbances within complex gene regulatory networks. Within these networks, certain genes hold pivotal roles, minor changes in their expression can disrupt the function of numerous downstream genes.These high-impact genes are often introduced as potential biomarkers or therapeutic targets and hold significant importance in precision medicine.

In this study, we constructed cancer-specific gene networks by selecting differentially expressed genes (DEGs) as the initial network nodes. The distribution of these DEGs was assessed using the Shapiro-Wilk test and QQ-plots, revealing that none followed a normal distribution. Subsequently, four correlation metrics were employed to identify gene-gene associations, and their similarities and differences were visualized through statistical plots and summary analyses. High-degree nodes (network hubs) were considered as potential pivotal genes in each cancer gene network. These hubs were then evaluated for their association with patient overall survival. Our results demonstrate that Spearman-based correlation networks were significantly more effective at selecting survival-related hub genes compared to other correlation methods. This advantage is likely due to Spearman’s ability to capture both linear and non-linear relationships while maintaining robustness against outliers.

To further validate the biological relevance of the detected associations, gene set enrichment analysis was conducted. The findings showed that hubs derived from Spearman-based networks enriched more biologically meaningful terms, both in quantity and quality, suggesting a higher functional relevance.

Despite these promising findings, First, this study focused solely on transcriptomic data, and integrating multi-omics layers (e.g., proteomics, epigenomics) may provide a more comprehensive understanding of gene regulatory mechanisms. Furthermore, the study used static gene expression datasets; however, gene networks are inherently dynamic. Time-series or single-cell data could better capture the temporal dynamics and heterogeneity of cancer cells. Third, although survival analysis was performed, functional validation of identified hub genes through in vitro or in vivo experiments remains necessary to confirm their biological and clinical relevance.

Future studies could aim to incorporate longitudinal or single-cell datasets, and explore machine learning-based network inference methods to complement correlation-based approaches. Additionally, expanding the framework to different cancer types and integrating clinical metadata could enhance the generalizability and translational potential of the findings.

## Material and Methods

### Data Collection

We evaluated the performance of each correlation method on four different RNA-seq datasets. The datasets were gene expression profiles of solid tissues for stomach adenocarcinoma (STAD), Bladder urothelial carcinoma (BLCA), Kidney renal clear cell carcinoma (KIRC) and Colon adenocarcinoma (COAD) which were downloaded from The Cancer Genome Atlas (TCGA, https://portal.gdc.cancer.gov) using TCGABiolinks package in R [16]. Since the research aim was the selection of survival-related genes, the cancers with lower survival rates were selected. The number of cancer samples and their survival information are depicted in Table 4.

**Table 4.**
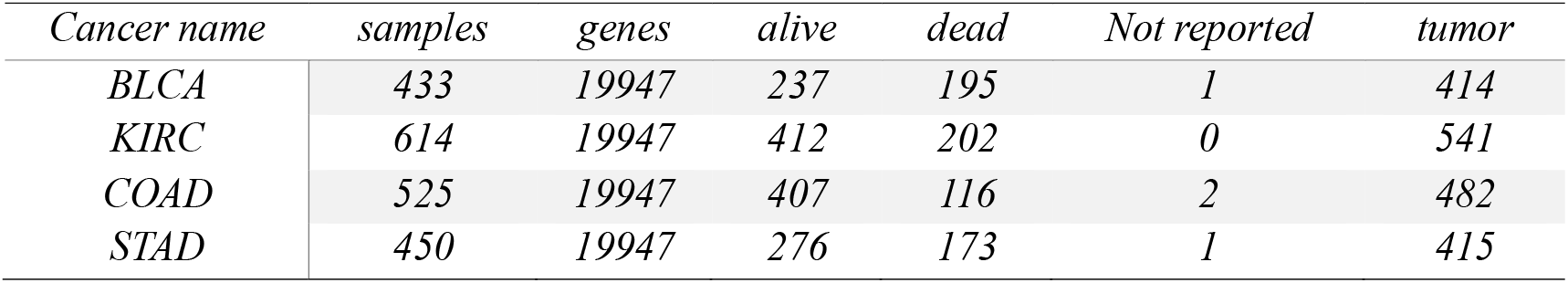
TCGA data characteristics.

| Cancer name | samples | genes | alive | dead | Not reported | tumor |
| --- | --- | --- | --- | --- | --- | --- |
| BLCA | 433 | 19947 | 237 | 195 | 1 | 414 |
| KIRC | 614 | 19947 | 412 | 202 | 0 | 541 |
| COAD | 525 | 19947 | 407 | 116 | 2 | 482 |
| STAD | 450 | 19947 | 276 | 173 | 1 | 415 |

Gene expressions were measured through mapping RNA-Seq by Expectation Maximization (RSEM). We first removed genes with RSEM expression values of 0 in all samples and then log 2 -transformed the expression levels.

### Normalization and differential expression

Expression profiles were normalized by the TCGAanalyze_Normalization function of TCGAbiolinks package in R. EdgeR package was used to identify the differentially expressed genes between the normal and tumor samples. Most significant differentially expressed genes with FDR < 0.05 and |logFC| > 1.5 were selected as initial nodes of cancer gene networks. According to this threshold, 1511 genes were selected as STAD, 2768 as BLCA, 2222 as KIRC, and 2303 as COAD initial networks’ nodes.

### Pearson’s correlation

Pearson’s correlation [17, 18] is a measure of linear dependence between two variables. It is the ratio between the covariance of two data sets and the product of their standard deviations.

Let 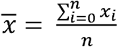 and 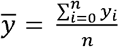 be the means of X and Y, respectively. Then, the Pearson’s correlation coefficient, **ρpearson** is defined as follows:

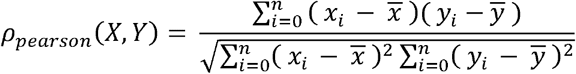

Pearson’s correlation is essentially a normalized measurement of the covariance, such that the result always has a value between −1 and 1. Pearson’s correlation is 1, in the case of positive (increasing) perfect linear dependence, and is -1 in the case of negative (decreasing) perfect linear dependence. The Pearson’s correlation is 0, in the case of linearly independent random variables, and in the case of imperfect linear dependence, -1< ρ_Pearson_ < 1. Sometimes, there is a misinterpretation of the last two cases because it is usually assumed that noncorrelated X and Y mean independent variables, whereas they may have nonlinear associations that Pearson’s correlation is not able to identify. The Python function for Pearson’s correlation is **corr** function with parameter (method **=** ‘Pearson’). Although default value for method in corr function is Pearson. You can also calculate Pearson’s correlation by **cor.test** function from **stats** package in R.

### Spearman’s correlation

Spearman’s rank correlation or simply Spearman’s correlation [19], unlike Pearson’s correlation, does not require assumptions of linearity in the relationship between variables, nor should the variables be measured at interval scales, since it can be used for ordinal variables [20]. For sample size n, the ranks for pair (x_i_, y_i_) are R[x_i_], R[y_i_], and ρ_Spearman_ is computed as follows:

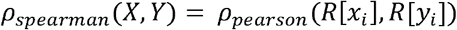

Only when all ranks are distinct integers (no ties), it can be computed as:

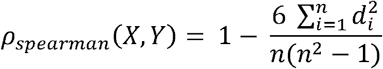

The Python function for Spearman’s correlation is **corr** function with parameter (method **=** ‘spearman’). You can also calculate Spearman’s correlation by **cor.test** function with parameter (method = ‘spearman’) from stats package in R.

### Kendall rank correlation

The Kendall rank correlation coefficient or simply Kendall’s correlation [21] is used to measure the ordinal association between two measured quantities. Kendall’s correlation is defined as follows:

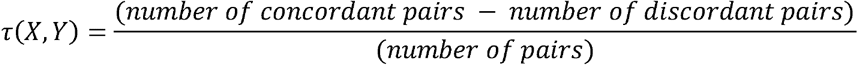

Where concordant reveals the agreement of elements ranks: that is x_i_ > and x_i_ > or in another scenario x_i_ < and y_i_ < . A pair of elements are discordant, if (x_i_ > and x_i_ < or y_i_ < and y_i_ >) . if x_i_ = x_i_ and y_i_ = y_i_, the pair is neither concordant nor discordant. “Number of pairs” declares the number of all cases in which we can select a pair of elements among n sample, which is equal to 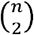.

Kendall’s correlation between two variables is high when elements have a similar rank between the two variables and low when elements have a dissimilar rank between the two variables. The Python function for Kendall’s correlation is corr function with parameter (method = ‘kendall’). You can also calculate Spearman’s correlation by cor.test function with parameter (method = ‘Kendall’) from stats package in R.

### Distance Correlation

Distance correlation or Distance covariance is a measure of dependence between two paired variables based on energy distances (a statistical distance between probability distributions) [22]. Let v_n_ (X, Y) be the empirical covariance, which is a non-negative number defined as follows:

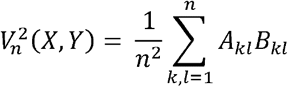

Where, A_kl_ and define as 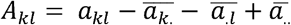 and 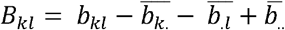 that 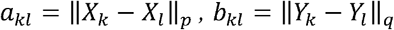 with k,l = 1, …, n. Again, for a random variable X, V_n_ (X) is non-negative and defines as:

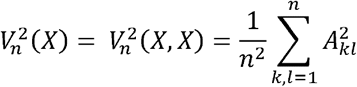

Eventually, the empirical correlation R_n_(X,Y)is the square root of bellow variable:

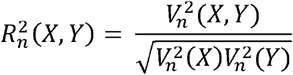

We calculate Distance correlation by **distance_correlation** from dcor package in Python. You can also calculate Distance correlation by dcor.test function from the energy package in R.

### Network construction by correlation methods

A network provides a natural representation of a complex cellular system in which nodes illustrate the molecular components and edges illustrate the biological relationships among these nodes(genes). After gene selections with significant differential expression as initial nodes for each cancer gene network. The four correlation methods were applied to define associations between genes (nodes). The four networks were built for each cancer based on specific correlation metrics. Two genes were associated in the network if their expression correlation was higher than 0.65. In this step, we just used tumor samples and omitted normal ones because they may be treated as noise and unreasonably increase the correlation.

An important development in understanding the cellular network architecture was the finding that most networks within the cell approximate a scale-free topology. This means a few nodes with a large number of links, often called hubs, hold the nodes together, and these hubs fundamentally determine the network’s behavior [1,23,24].

We selected the high-degree nodes as network hubs for each cancer gene network since these genes could have a critical role in cancer initiation, progression, or treatment.

### Survival analysis

Kaplan–Meier and log-rank tests were used to determine the association between the gene expressions and overall survival. Among those genes with significant differential expression, the samples were labeled as “Low count” and “High count” according to their expression compared to the median gene expression among all cancer samples. Statistical significance was also set at p-value < 0.05. Finally, those genes whose expression among different samples was significantly associated with OS were selected. The survival analysis was done by survminer package in R. Finally, we found 207 genes related to OS in STAD, 643 in BLCA, 914 in KIRC, and 133 in COAD.

### Enrichment analysis

Gene set enrichment analysis is a bioinformatics technique that identifies the most over-represented biological processes or biological pathways in a list of genes compared to those that would be associated with them by chance. We applied g profiler (https://biit.cs.ut.ee/gprofiler/gost) [25] for enrichment analysis.

## Abbreviation

STAD: Stomach Adenocarcinoma
COAD: Colon Adenocarcinoma
BLCA: Bladder urothelial carcinoma
KIRC: kidney Renal Clear Cell Carcinoma
GO: Gene Ontology
OS: Overall Survival
QQ-plot: Quantile-Quantile plot
MF: Molecular Function
BP: Biological process
CC: Cellular Component

## Declarations

### Ethics approval

“Not applicable”

### Informed Consent

“Not applicable”

### Availability of data and material

The gene expression profiles were downloaded from TCGA (https://portal.gdc.cancer.gov) and the source code is available in https://github.com/Amirhosein-JVD/Correlation-Analysis. The rest of the information included within the article and its additional files.

### Authors’ contributions

HR conceived the research. HR, AJ and CE designed the research. HR and AJ implemented the research. HR wrote the manuscript. All authors read and approved the final manuscript.

## Acknowledgements

The authors would like to thank editors and reviewers for their insightful suggestions and careful reading of manuscript

## Supplementary Information

Supplementary File 1. Lists of genes that were differentially expressed among normal and tumor samples for each cancer.

Supplementary File 2. List of survival-related gens for each cancer (Results of Kaplan–Meier and log-rank tests) Supplementary File 3. Results of g:Profiler enrichment analysis for each cancer.

Supplementary File 4. Lists of enriched GO_terms that are associated with cancer hallmarks.

